# A Synthetic Mirtron Platform Enables Stable and Robust Splicing-Dependent Gene Silencing in Plants

**DOI:** 10.64898/2026.03.04.709674

**Authors:** Nitsan Lugassi, Bolaji Jay Isaac, Shoval Nitsani, Ben Spitzer-Rimon, Kinneret Shefer

## Abstract

Post-transcriptional gene silencing (PTGS) is widely used for gene function studies and crop improvement; however, conventional transgene-based RNAi and artificial microRNA (amiRNA) approaches are often subject to transgene self-silencing, epigenetic inactivation, viral suppressors of RNA silencing, and regulatory complexity. Here, we establish for the first time a synthetic mirtron platform in plants that defines a splice-gated, non-canonical PTGS architecture, mechanistically distinct from existing RNAi strategies. We demonstrate that precise intron splicing and lariat debranching are essential for target gene silencing, directly coupling pre-mRNA splicing to small RNA-mediated regulation. An optimized mirtron mediates efficient, stable, and heritable silencing of *PHYTOENE DESATURASE* in *Arabidopsis thaliana* and enables multiplex silencing of endogenous *AUXIN RESPONSE FACTORS* in potato, demonstrating applicability in a major crop species. Moreover, mirtron-mediated gene silencing remains highly effective in the presence of the viral suppressor P19, unlike canonical amiRNA-based silencing, highlighting its greater resistance to viral suppression and utility for host-induced gene silencing. Because mirtrons are embedded within endogenous introns, they can be co-expressed with host genes, inheriting their native spatial and temporal expression patterns. When precisely introduced, this architecture supports regulatory outcomes similar to those of gene-edited products that lack foreign DNA. Together, these findings define mirtrons as a compact, stable, and application-ready PTGS platform useful for studying gene regulation and crop biotechnology.

## Introduction

Global food security has emerged as a major challenge as population growth continues alongside climate change, imposing increasing constraints on agricultural production systems. Together, these pressures intensify the demand for innovative crop improvement strategies that deliver enhanced agronomic performance, increased resilience, higher, more stable yields, and improved nutritional quality (Charles et al., 2010; Hickey et al., 2019). Traditional breeding methods have long provided, and continue to provide, solutions to agricultural challenges. However, while effective over time, these approaches are often slow and, in many cases, limited by evolutionary bottlenecks that limit available genetic variation, as well as by genetic linkage of traits that constrains the independent selection of desirable characteristics (Allaby et al., 2019; Hickey et al., 2019; Pixley et al., 2022;) Consequently, such limitations can hinder the ability to keep pace with the rapid development of solutions required to address current agricultural challenges. Genetically modified (GM) plants offer a more immediate approach by enabling the direct introduction of defined genetic traits, thereby bypassing constraints imposed by limited natural variation and genetic linkage (Hickey et al., 2019). More recently, genome-editing technologies have further expanded the plant biotechnology toolbox, enabling targeted and precise modification of endogenous genes (Pixley et al., 2022). Nevertheless, GM approaches face practical limitations, most notably regulatory constraints and the risk of transgene silencing. Genome editing strategies, while alleviating some of these barriers, continue to face significant biological challenges, particularly when targeting essential genes, achieving efficient, uniform modification across polyploid or highly heterozygous genomes, and dealing with redundant gene families. (He and Krainer, 2021; Turnbull et al., 2021; Zhang et al., 2022; Cardi et al., 2023)

Endogenous small RNA pathways in plants, including microRNA (miRNA) and small interfering RNA (siRNA) pathways, provide the mechanistic basis for post-transcriptional gene silencing and have been widely harnessed for crop improvement through RNA interference (RNAi) and artificial microRNA (amiRNA)–based approaches (Guo et al., 2016). Hairpin, inverted-repeat RNAi, and amiRNA strategies enable efficient suppression of individual genes and gene families and have contributed to gene function studies and offer opportunities for the improvement of diverse agronomic traits, including yield-related characteristics, quality attributes, secondary metabolism, and stress tolerance (Schwab et al., 2006; Carbonell et al., 2014; Koch and Kogel, 2014; Saurabh et al., 2014; Kamthan et al., 2015; Chen et al., 2021; Ullah et al., 2022; Dharshini et al., 2025; Kumar et al., 2025). In addition, these approaches have been extended to crop protection through host-induced gene silencing (HIGS), in which plant-expressed double-stranded RNAs target essential genes in pests and pathogens (Nowara et al., 2010; Koch and Kogel, 2014; Bramlett et al., 2020).

Despite these advantages, existing PTGS-based approaches can be limited by variability in silencing efficiency and stability that depend on transgene silencing, epigenetic modification, expression levels, selected transgene promoter and terminator, and the genomic context of the transgene (Matzke and Matzke, 1998; F de Felippes et al., 2020; Feng et al., 2022; Zhang et al., 2022; Koo et al., 2025; Roca Paixao and Déléris, 2025). Moreover, RNAi-based gene silencing is susceptible to suppression by viral silencing suppressors that directly antagonize host RNA silencing pathways, thereby enhancing viral accumulation. A prominent example is the tombusvirus suppressor p19, which binds and sequesters 21-nt siRNA duplexes, preventing their loading into the RNA-induced silencing complex and thereby inhibiting systemic post-transcriptional gene silencing and counteracting RNAi-mediated defenses (Lopez-Gomollon and Baulcombe, 2022; Vaucheret and Voinnet, 2024). These limitations highlight the need for alternative small RNA architectures that engage RNA silencing pathways through distinct biogenesis routes.

In plants, most miRNAs are intergenic and expressed as independent transcription units, whereas in animals a substantial fraction of miRNAs is intronic and co-transcribed with their host genes (Brown et al., 2008; Shomron and Levy, 2009; Voinnet, 2009). In a limited subset of cases in animals, a distinct class of intronic miRNAs, termed mirtrons, bypasses canonical Drosha processing (Ruby et al., 2007; Westholm and Lai, 2011). Canonical miRNA precursors and mirtrons differ in their structural organization and boundary definition, which leads to differences in their processing pathways (Ruby et al., 2007; Westholm and Lai, 2011). Canonical miRNAs are transcribed as long primary transcripts (pri-miRNAs) containing an internally defined stem–loop embedded within extended flanking sequences; this structure is recognized and cleaved in the nucleus by the Drosha–DGCR8 microprocessor complex to release the precursor hairpin. In contrast, mirtrons originate from short introns whose boundaries correspond precisely to splice donor and acceptor sites(Ruby et al., 2007; Westholm and Lai, 2011). Following spliceosomal excision and lariat debranching, the entire intron folds into a pre-miRNA-like hairpin lacking extended flanking regions, with its ends defined by splice junctions rather than by endonucleolytic cropping. Consequently, mirtron biogenesis bypasses Drosha processing and instead directly couples pre-mRNA splicing to Dicer-mediated maturation, ultimately producing ∼21–23 nucleotide small RNAs that are incorporated into ARGONAUTE-containing silencing complexes(Ruby et al., 2007; Westholm and Lai, 2011).

In plants, no functional native mirtrons have been identified, nor have artificial mirtrons previously been used to induce gene silencing; however, intron-embedded amiRNA designs have been shown to induce gene silencing, demonstrating that the plant RNA processing machinery can support intron-associated small RNA biogenesis (Shapulatov et al., 2018). In these systems, however, the intron primarily serves as a transcriptional or structural context for a canonical pri-miRNA hairpin, and the miRNA maturation proceeds through the standard DCL1 pathway. In the current study, we establish a synthetic mirtron platform as a functional, splicing-dependent gene silencing system in plants. We show that precise intron splicing and lariat debranching are essential for small RNA production and target repression, allowing efficient, stable, and heritable silencing without transgene self-silencing. Using this platform, we demonstrate tunable, multiplexed gene silencing in both Arabidopsis and potato, as well as reduced susceptibility to P19-Mediated Viral silencing suppression. Together, these findings demonstrate mirtrons as a mechanistically distinct and application-ready PTGS architecture that expands the toolkit for gene regulation research and crop biotechnology.

## Results

### Establishment of splicing-dependent silencing of *PDS* via a mirtron-based system in *Arabidopsis*

Functional mirtrons are well established in animals but have not been experimentally demonstrated in plants. Mirtrons depend on precise intron excision, correct lariat formation, and proper release of a hairpin precursor following splicing. To establish a mirtron-based silencing system in plants, we designed a series of synthetic mirtron candidates that varied in predicted splicing efficiency. To directly compare silencing efficacy, all constructs contained the same 21-nt sequence targeting the coding region of *Arabidopsis thaliana PHYTOENE DESATURASE* (*PDS*; nucleotides 21–41 of the CDS; Figure 1A). Each mirtron–PDS cassette was embedded as an intron within the *GFP* coding sequence and expressed under the constitutive CaMV 35S promoter (Figure 1A). In this configuration, successful intron splicing restores GFP fluorescence, whereas effective *PDS* silencing produces the characteristic albino phenotype. A predicted functional mirtron (construct 94) was designed with canonical 5′ splice donor and 3′ splice acceptor sites and intronic sequences (Figure 1A). The engineered sequence was predicted to form a stable hairpin (Mfold; Zuker, 2003). Following stable transformation, transgenic *Arabidopsis* plants carrying construct 94 (hereafter, 35S::GFP-MIRT_PDS_) exhibited a strong albino phenotype, along with robust GFP fluorescence (Figure 1B–C), indicating efficient intron splicing and effective *PDS* silencing. In contrast, transgenic *Arabidopsis* plants carrying a control construct in which the mirtron was replaced with a native *Arabidopsis* intron (the 15th intron of AT3G60950; 35S::GFP-INT) exhibited strong GFP fluorescence but retained normal green pigmentation, as wild-type plants (Figure 1D–G). Sequencing of spliced cDNA products derived from 35S::GFP-MIRT_PDS_ and 35S::GFP-INT confirmed accurate intron excision and precise splice junction formation in the functional mirtron construct (Supplementary Figure 1). These results confirm that restoration of GFP expression alone is insufficient to induce *PDS* silencing and that silencing is specifically mediated by the engineered mirtron sequence.

**Figure 1.**
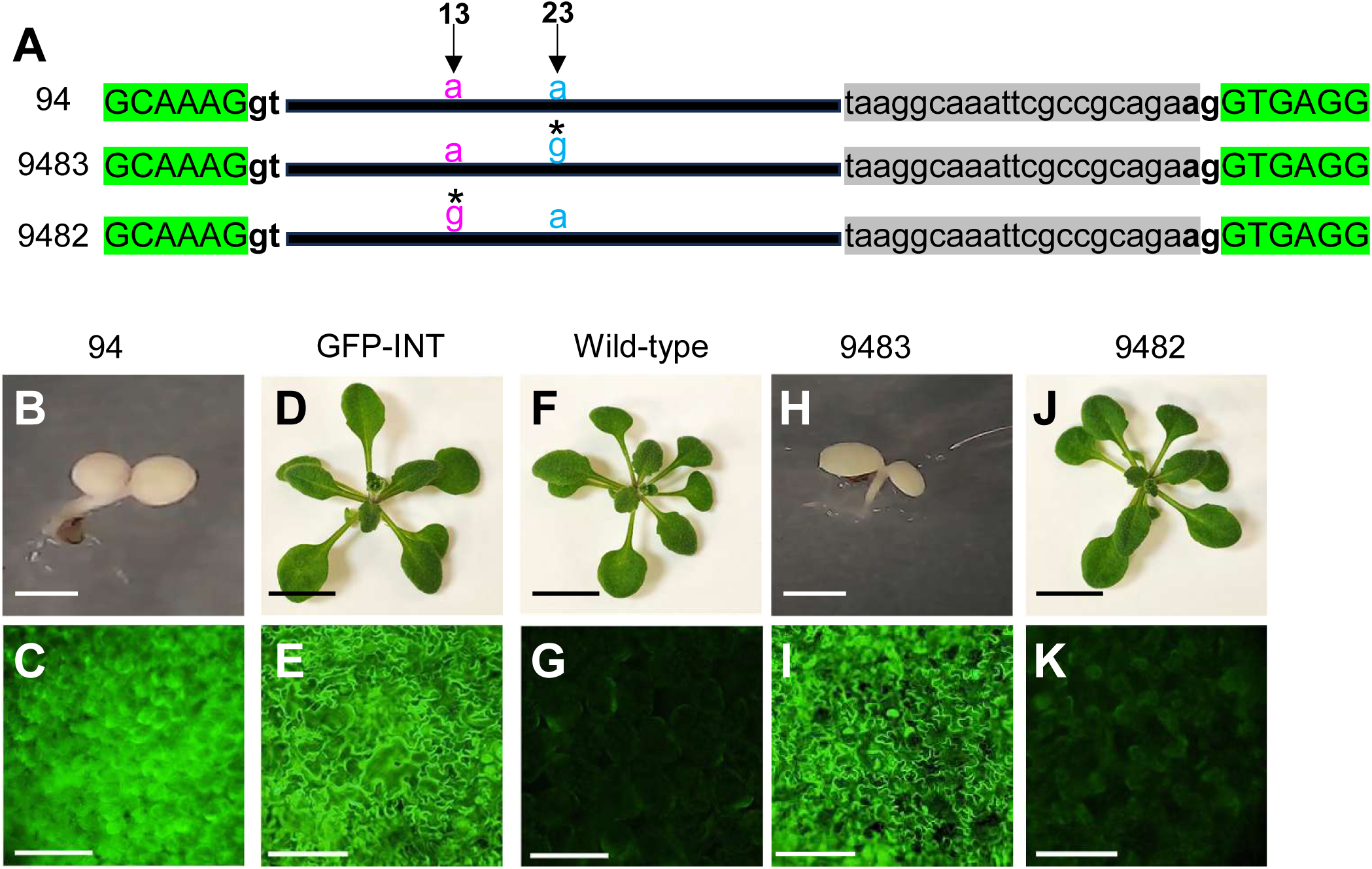
Splicing-dependent mirtron-mediated silencing of *PDS* in *Arabidopsis*. (A) Schematic representation of synthetic mirtron constructs embedded within the *GFP* coding sequence. Construct numbers are indicated on the left. Capital letters in green represent the *GFP* CDS. Canonical splice donor (GT) and acceptor (AG) sites are shown in bold. The mirtron–*PDS* seed sequence is indicated in gray. Asterisks (*) represent nucleotide substitutions relative to construct 94. Magenta and blue nucleotides indicate candidate branch-point adenosines, with their positions relative to the 5′ splice site shown above. The black line represents the remaining intronic sequence. (B–K) Representative T_1_ transgenic plants expressing the indicated constructs, and wild-type control plants. (B–C) 35S::GFP-MIRT_PDS_ (construct 94). (D–E) Native intron control (GFP-INT). (F–G) Wild-type plants. (H–I) Construct 9483. (J–K) Construct 9482 (35S::GFP-MIRT_PDS_-NSC). (B, D, F, H, J) Bright-field images. (C, E, G, I, K) Corresponding GFP fluorescence images. Scale bars represent 1 cm in (D, F, J), 2 mm in (B, H), and 100 μm in (C, E, G, I, K).

To identify the functional branch-point A required for the splicing of 35S::GFP-MIRT_PDS_ and to determine whether splicing is essential for silencing of the engineered mirtron, two additional constructs, nearly identical to 35S::GFP-MIRT_PDS_, were generated in which candidate intronic adenines were mutated to guanine. Construct 9483 carries an A to G substitution at position 23 downstream to the 5′ splice site, whereas construct 9482 carries an A to G substitution at position 13 downstream to the 5′ splice site (Figure 1A). Transgenic *Arabidopsis* plants carrying construct 9483 retained both GFP fluorescence and the albino phenotype, indicating that this mutation did not disrupt intron splicing (Figure 1H–I). In contrast, plants transformed with construct 9482 showed neither GFP fluorescence nor albino phenotype, demonstrating loss of splicing activity and consequent loss of silencing (Figure 1J–K). These findings identify the adenine at position 13 relative to the 5′ splice site as the functional branch point in the design of the 35S::GFP-MIRT_PDS_ mirtron construct. Construct 9482 was therefore designated as a non-splicing control (NSC; 35S::GFP-MIRT_PDS_-NSC). Together, these results demonstrate that efficient and precise intron splicing is essential for mirtron-mediated gene silencing in plants. The 35S::GFP-MIRT_PDS_ construct fulfills the defining functional criteria of a mirtron, providing direct experimental evidence that mirtron-based architectures can operate as a regulated RNA silencing platform in plants.

### 35S::GFP-MIRTPDS transgenic plants exhibit high-efficiency and stable PDS silencing without self-silencing

To assess the efficiency of mirtron-induced gene silencing, we quantified the proportion of albino plants among 35S::GFP-MIRT_PDS_ transgenic lines. Across T_1_ progeny derived from five independent T_0_ transformation events, 155 lines were scored for the presence or absence of the albino phenotype. A high proportion of lines (87% ± 3.2% SE) exhibited a complete albino phenotype, indicating relatively highly efficient silencing of *PDS*. To evaluate the durability and temporal stability of the silencing phenotype, we examined whether mirtron-mediated silencing is susceptible to transgene self-silencing, a known limitation of conventional RNAi approaches in plants (Dong et al., 2011; Lu et al., 2015; Zhang et al., 2022). Twenty independent albino T1 lines were transferred to non-selective medium (Figure 2A) and monitored for up to three months. All plants retained a stable albino phenotype with no evidence of phenotypic reversion (Figure 2A–B). This experiment was independently repeated three times (total n = 60 lines), yielding consistent results.

**Figure 2.**
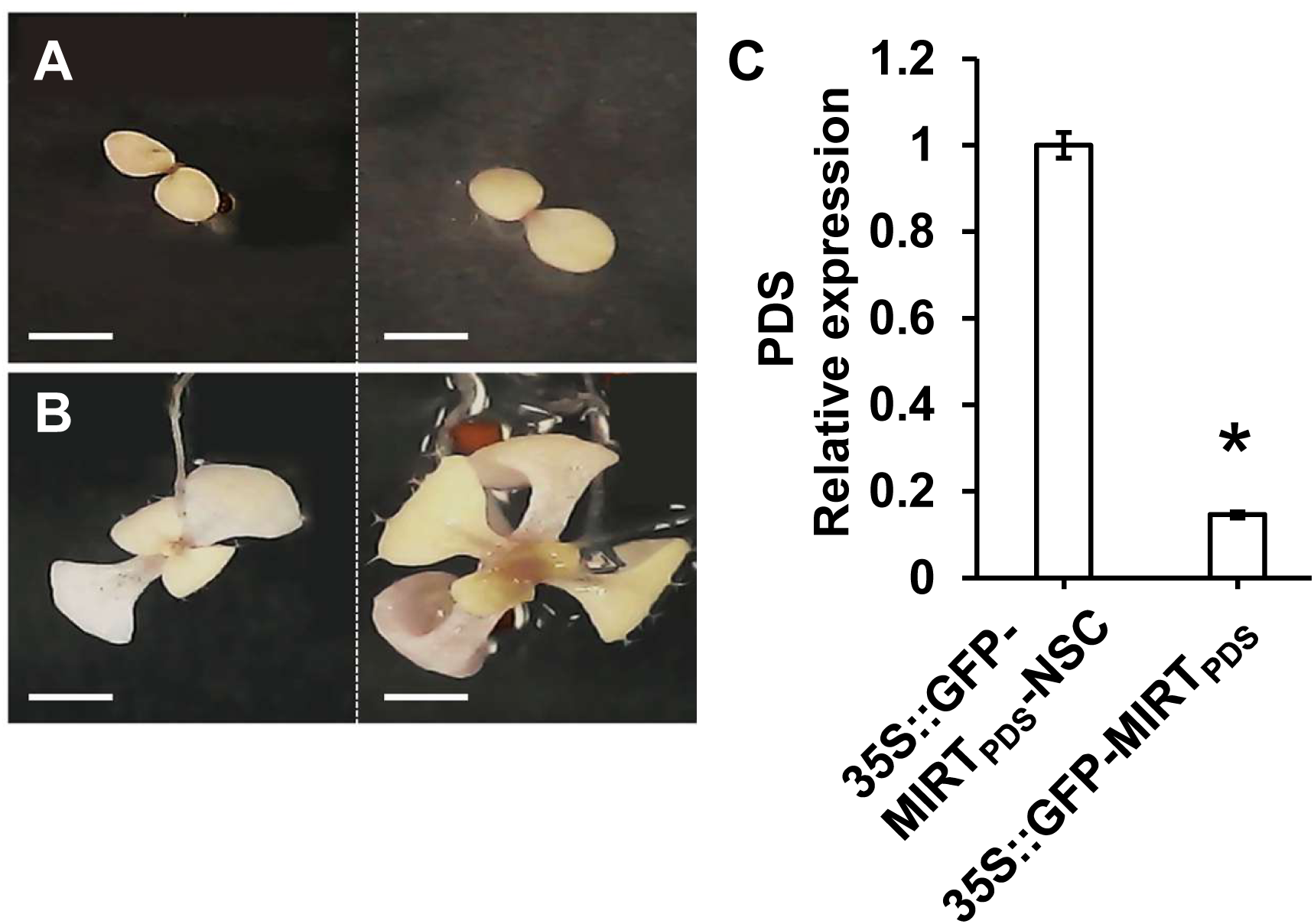
Temporal stability and durability of mirtron-mediated *PDS* silencing. (A) Representative of two albino T1 plants expressing 35S::GFP-MIRT_PDS_ following transfer from selective conditions to non-selective medium. (B) The same plants are shown after three months of growth on non-selective medium. Scale bars represent 2 mm. (C) Quantitative real-time RT–PCR analysis of *PDS* transcript levels in 35S::GFP-MIRT_PDS_ plants relative to 35S::GFP-MIRT_PDS_-NSC control plants. Bars represent mean ± standard error of 4 biological replicates; asterisk indicates a statistically significant difference (*P* < 0.01, Student’s *t*-test).

To molecularly validate mirtron-induced *PDS* silencing, quantitative real-time RT–PCR analysis was performed. *PDS* transcript levels were significantly reduced in 35S::GFP-MIRT_PDS_ albino transgenic plants compared to 35S::GFP-MIRT_PDS_-NSC transgenic plants (Figure 2C), confirming robust transcript depletion. Taken together, these results demonstrate that 35S::GFP-MIRT_PDS_ mediates high-efficiency and durable *PDS* silencing, without evidence of transgene self-silencing, a phenomenon commonly associated with conventional RNAi constructs.

### Tunable and heritable mirtron-mediated *PDS* silencing

To determine whether the extent of mirtron-induced gene silencing could be modulated through alterations in the seed sequence, we introduced a single-nucleotide substitution into the seed region of the *PDS*-targeting mirtron, generating the 35S::GFP-MIRT_PDS_(A18T) construct (Figure 3A). This A-to-T substitution at position 18 within the seed sequence was designed to reduce target complementarity and thereby attenuate silencing efficiency. Multiple independent T_1_ transgenic plants expressing 35S::GFP-MIRT_PDS_(A18T) exhibited intermediate albino phenotypes that were clearly distinct from the complete albinism observed in plants expressing the original 35S::GFP-MIRT_PDS_ construct (Figure 3B–D). From these, two representative lines carrying a single T-DNA insertion and fixed to homozygosity were selected for further analysis of phenotypic inheritance. All T_4_ progeny derived from these lines consistently displayed the partial albino phenotype, demonstrating that the attenuated silencing effect was stably maintained across generations. To quantify the extent of *PDS* silencing in the partial albino-T4 plants, chlorophyll content (Figure 3E) and *PDS* transcript levels (Figure 3F) were assessed. Both analyses revealed reduced *PDS* expression and chlorophyll accumulation in 35S::GFP-MIRT_PDS_(A18T) plants, relative to the 35S::GFP-MIRT_PDS_-NSC transgenic plants, but substantially higher levels than those observed in plants transgenic for 35S::GFP-MIRT_PDS_ (Figure 3E-F), consistent with an intermediate silencing phenotype. Together, these results demonstrate that the magnitude of mirtron-mediated gene silencing can be fine-tuned through seed-sequence modifications and that mirtron-based silencing is heritable and can be stably maintained across multiple generations.

**Figure 3.**
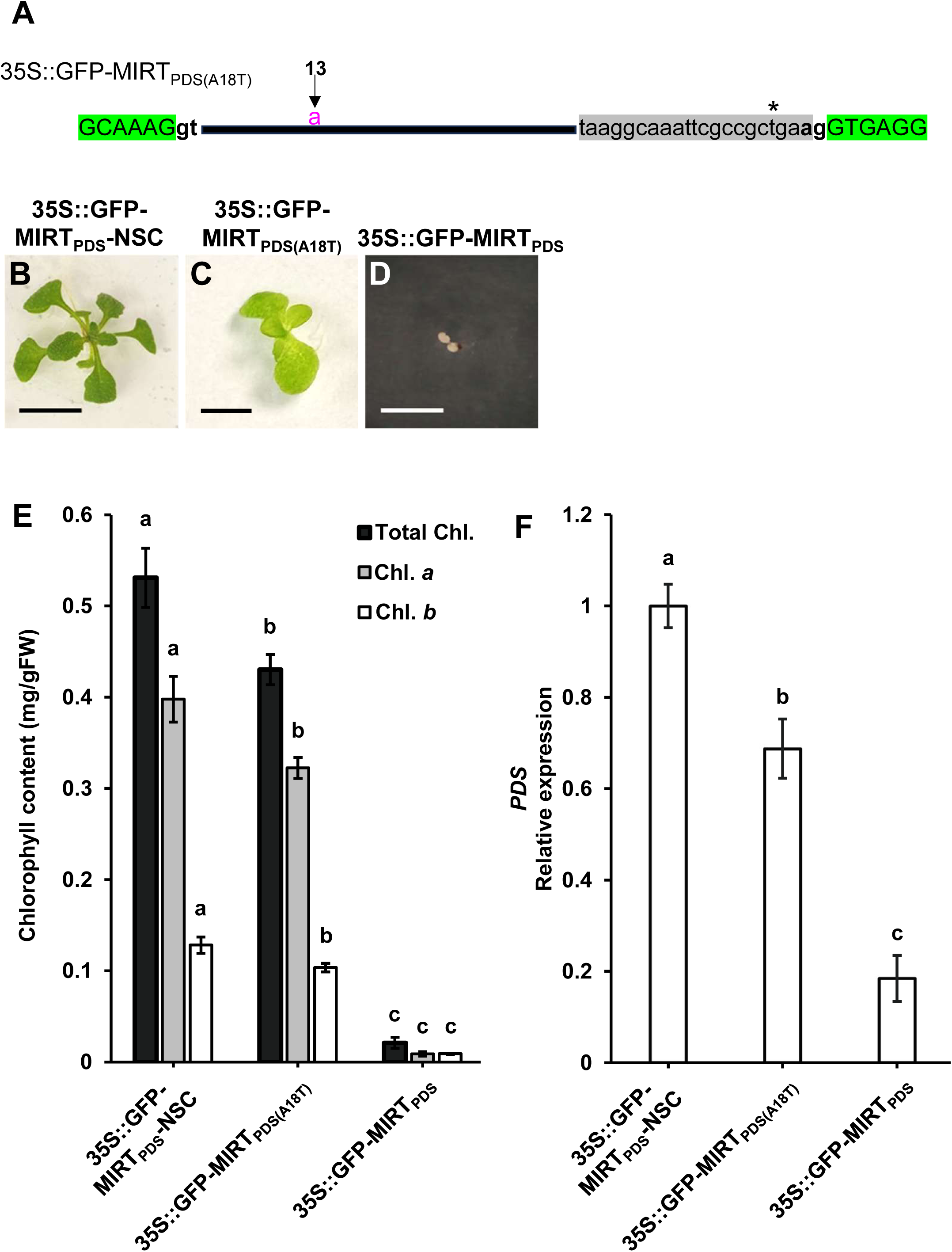
Seed sequence modification enables tunable mirtron-mediated silencing. (A) Schematic representation of the synthetic mirtron 35S::GFP-MIRT_PDS(A18T)_ construct embedded within the *GFP* coding sequence. Capital letters in green represent the *GFP* CDS. Canonical splice donor (GT) and acceptor (AG) sites are shown in bold. The mirtron–PDS seed sequence is indicated in gray. Asterisks (*) represent A-to-T nucleotide substitutions. Magenta nucleotide indicates branch-point adenosine, with its positions relative to the 5′ splice site shown above. The black line represents the remaining intronic sequence. (B–D) Representative T1 plants transgenic for: (B) 35S::GFP-MIRT_PDS_-NSC, (C) 35S::GFP-MIRT_PDS(A18T)_, and (D) 35S::GFP-MIRT_PDS_. Scale bars represent 1 cm. (E) Chlorophyll *a* (Chl. *a*), chlorophyll *b* (Chl. *b*), total chlorophyll content (Total Chl) in transgenic for 35S::GFP-MIRT_PDS_-NSC, 35S::GFP-MIRT_PDS(A18T)_, and 35S::GFP-MIRT_PDS_. (F) Quantitative real-time RT–PCR analysis of *PDS* transcript levels normalized to *TUB2.* For panels (E) and (F), analyses were performed on T4 lines for all constructs except 35S::GFP-MIRT_PDS_, for which independent T1 lines were used due to the inability of fully albino plants to reach reproductive maturity; bars represent mean ± standard error from three to six biological replicates. Statistical analysis was performed using one-way analysis of variance (ANOVA) followed by Tukey’s post hoc test, conducted independently for each assay. Different letters indicate statistically significant differences of (*P* < 0.05) for (E) and (*P* < 0.01) for (F).

### Mirtron-mediated silencing of multiple targets in potato

To evaluate the ability of mirtron technology to simultaneously target multiple genes and to assess its activity in a crop species, we targeted the potato *AUXIN RESPONSE FACTOR* (*StARF*) genes: *StARF10 and StARF17,* by embedding the *miR160* seed sequence into a mirtron–GFP construct driven by the CaMV 35S promoter (35S::GFP-MIRT_160_). Three independent transgenic lines (L22-9, L22-20, and L22-28) exhibited reproducible developmental abnormalities, including dwarfism and leaf curvature (Figure 4A-H). Quantitative real-time RT-PCR analysis revealed significant downregulation of both *StARF10* and *StARF17* transcripts relative to non-transgenic control plants in all three lines (Figure 4I-J). These results indicate that a single mirtron construct can mediate coordinated and effective silencing of multiple endogenous targets in potato, demonstrating the feasibility of mirtron-based gene silencing for functional studies of multi-gene families and for multiplex trait modulation in crop plants.

**Figure 4.**
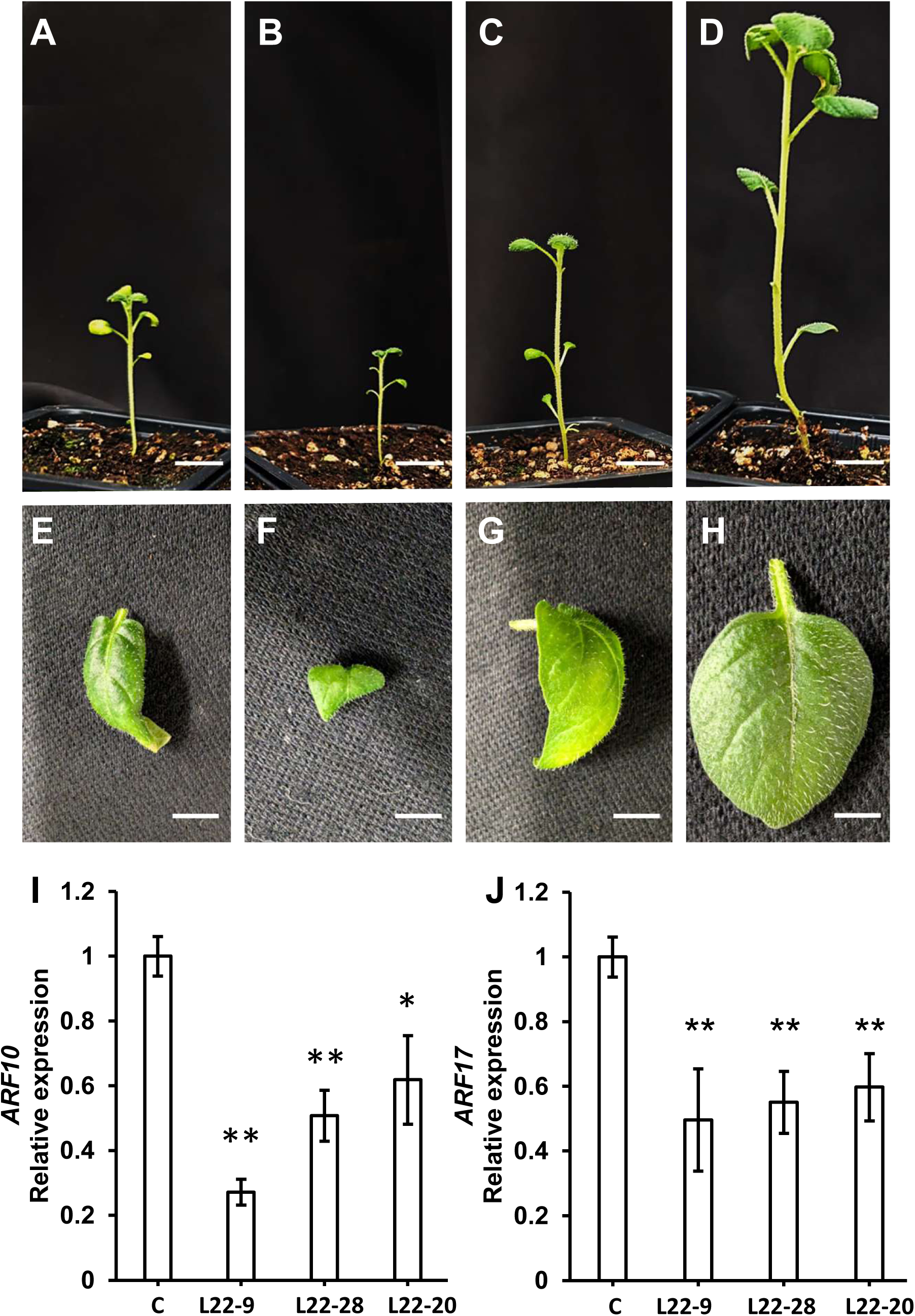
Mirtron-mediated silencing of multiple targets in potato using a single mirtron construct. (A–D) Representative whole-plant phenotypes and (E–H) corresponding leaf morphology of potato plants expressing 35S::GFP-MIRT_160_ in three independent transgenic lines. (A, E) line L22-9. (B, F) line L22-28. (C, G) L22-20. Non-transgenic control (D, H). Scale bars represent 1 cm in (A–D) and 0.5 cm in (E–H). (I–J) Quantitative real-time RT–PCR analysis of *ARF10* (I) and *ARF17* (J) transcript levels, normalized to *ElF3e*, in transgenic lines L22-9, L22-20, and L22-28 relative to non-transgenic control plants. Bars represent the mean ± standard error of at least three biological replicates. Significance of differences (\**P* < 0.05; \*\**P* < 0.01) between treatments and control was calculated using Dunnett’s method following analysis of variance (ANOVA) with the non-transgenic control as the reference.

### Mirtron-mediated silencing is resistant to viral silencing suppression

A major limitation of PTGS–based strategies for crop biotechnology and HIGS is their susceptibility to viral silencing suppressors (Lopez-Gomollon and Baulcombe, 2022; Vaucheret and Voinnet, 2024). To evaluate the sensitivity of mirtron-mediated silencing to viral suppression in plants, we examined *PDS* silencing in the presence of the *Tomato bushy stunt virus* silencing suppressor P19. *Arabidopsis* plants were transformed with four constructs: GFP-MIRT_PDS_ or amiRNA_PDS_ expressed under the *HIGH CONSTITUTIVE PROMOTER1* (*pCH1*) promoter (pPCH1::GFP-MIRT_PDS_ and pCH1::amiRNA_PDS_, respectively; Zhou et al., 2023), either alone or together with *P19* expressed from the CaMV 35S promoter (Figure 5A). The *pCH1* promoter was used to drive moderate constitutive expression of the silencing constructs, whereas P19 was expressed from the CaMV 35S promoter to ensure robust accumulation of the viral silencing suppressor. Both pCH1::GFP-MIRT_PDS_ and pCH1::amiRNA_PDS_ constructs contained the same *PDS* seed sequence. Nine days after germination, T1 lines displaying clear *PDS* silencing phenotypes were selected, transferred to fresh medium, and monitored for an additional eight days (Figure 5B-K). While 44% of plants transgenic for pCH1::amiRNA_PDS_ together with P19 exhibited partial recovery of growth and pigmentation, only ∼7% of plants transgenic for pCH1::GFP-MIRT_PDS_ together with P19 showed recovery, a frequency comparable to that observed in control plants transgenic for pCH1::GFP-MIRT_PDS_ or pCH1::amiRNA_PDS_ alone (Figure 5L). These results indicate that mirtron-mediated silencing remains highly effective despite the presence of the viral suppressor P19, whereas canonical amiRNA-based silencing is susceptible to suppression by the viral silencing suppressor, supporting the notion that mirtron-mediated PTGS is not subject to viral suppression.

**Figure 5.**
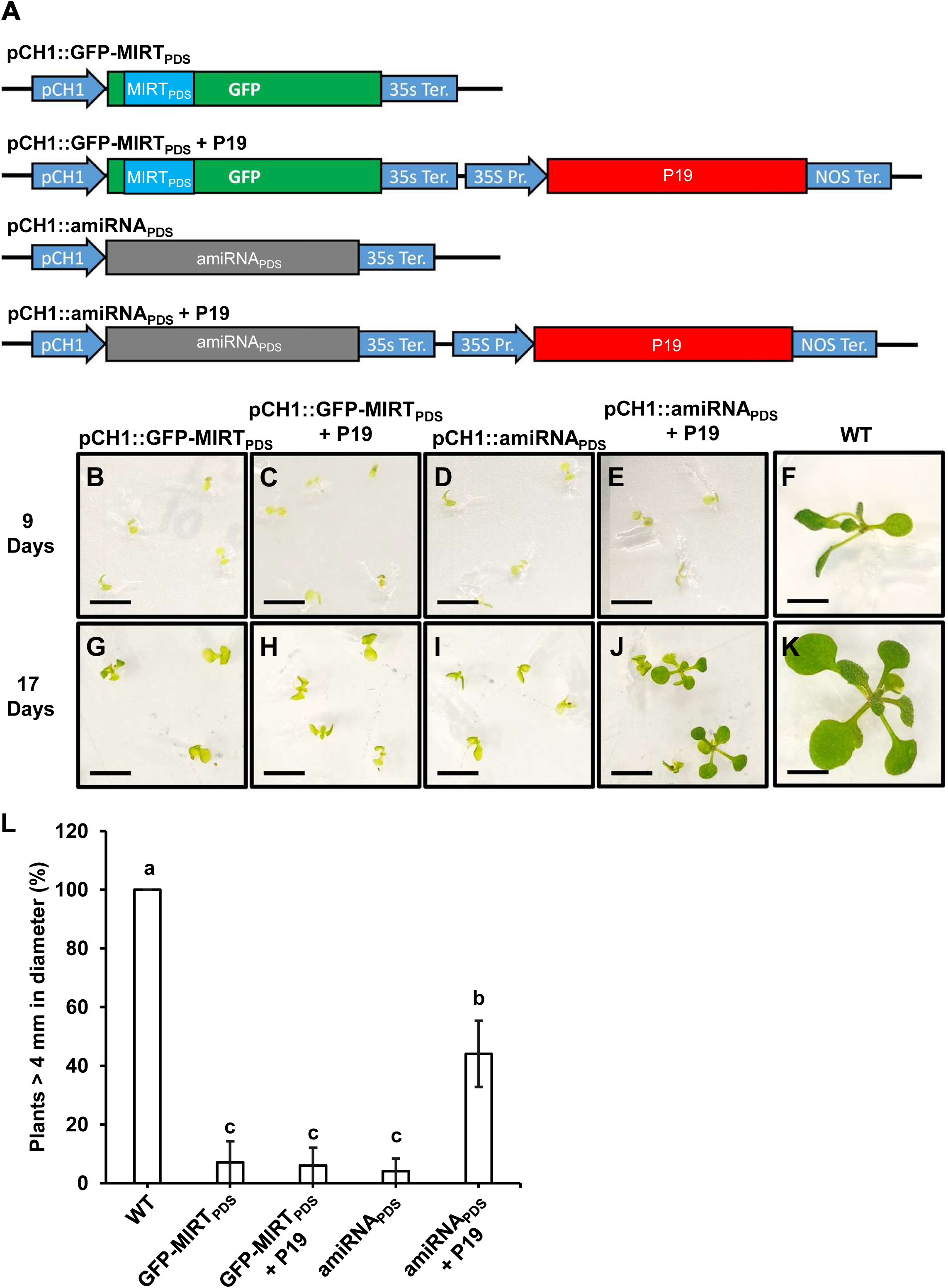
Reduced sensitivity of mirtron-mediated *PDS* silencing to P19 viral silencing suppression. (A) Schematic representation of expression constructs used for analysis of viral silencing suppressor sensitivity. (B–K) Representative T1 *Arabidopsis* seedlings nine days after germination, transgenic for pCH1::GFP-MIRT_PDS_ (B, G), pCH1::GFP-MIRT_PDS_ + P19 (C, H), pCH1::amiRNA_PDS_ (D, I), pCH1::amiRNA_PDS_ + P19 (E, J), or wild-type controls (F, K). Seedlings were transferred to fresh medium (B-F) and imaged after an additional eight days (G-K). Scale bars represent 0.5 cm. (L) Quantification of phenotypic recovery frequency in T1 plants expressing the indicated constructs. Bars represent the mean ± standard error of at least 26 independent lines. Statistical analysis was performed using one-way analysis of variance (ANOVA) followed by Tukey’s post hoc test. Different letters indicate statistically significant differences (P < 0.01). High constitutive promoter1 (pCH1); Cauliflower mosaic virus *35S* terminator (35S Ter.); Cauliflower mosaic virus *35S* promoter (35S Pr.); nopaline synthase terminator (NOS Ter.)

## Discussion

This study provides the first direct experimental demonstration that a synthetic mirtron-based architecture can function as an effective, spliceosome-dependent gene-silencing platform in plants. Although mirtrons are well-established in animals, their functional activity in plants has not been previously shown (Ruby et al., 2007; Westholm and Lai, 2011; Joshi et al., 2012; Formey et al., 2014; Shapulatov et al., 2018). Using rationally engineered intron-embedded hairpins, we demonstrate that efficient and precise intron splicing is required to generate an active silencing trigger in plants. The functional mirtron construct simultaneously restored GFP expression and induced robust *PDS* silencing, directly coupling spliceosome-mediated intron excision to sequence-specific post-transcriptional repression (Figure 1). The branch-point adenosine initiates lariat formation during spliceosome assembly and is essential for intron excision. In the 35S::GFP-MIRT_PDS_ construct, position 13 relative to the 5′ splice site was identified as the functional branch point. The mutation of this adenosine abolished both splicing and silencing (Figure 1). Optimization of future mirtron designs will require careful identification of the functional branch-point adenosine, as changes in the mirtron sequence or structure may alter its position. Together, these results establish that mirtron-mediated silencing in plants is strictly dependent on accurate spliceosomal processing and that small RNA biogenesis can be gated by intron excision.

This mirtron processing mechanism differs fundamentally from previously described intron-embedded amiRNA strategies in which introns primarily serve as an expression context for canonical pre-miRNA precursors (Shapulatov et al., 2018). In that system, miRNA-mediated silencing was not demonstrated to require accurate spliceosomal excision, and silencing activity persisted even when splicing was inefficient or aberrant. In contrast, synthetic mirtron-mediated silencing is strictly contingent upon precise intron excision and subsequent lariat debranching, as the functional hairpin is generated only after correct spliceosome-mediated processing (Figure 1; Ruby et al., 2007; Westholm and Lai, 2011). Thus, rather than embedding a conventional miRNA within an intron, the synthetic mirtron platform directly couples small RNA biogenesis to accurate spliceosome activity, establishing a genuinely splice-gated and sequence-programmable mechanism for post-transcriptional gene silencing in plants.

A key advantage of mirtron-mediated silencing observed here is its high stability and heritable maintenance, with no evidence of self-silencing of the transgene. The majority of 35S::GFP-MIRT_PDS_ transgenic lines (87%) displayed robust silencing, and the albino phenotype remained stable throughout the entire experimental period, which extended for at least three months (Figure 2). In advanced generations (T_4_), lines carrying the *PDS* seed sequence–mutated mirtron variant consistently maintained an intermediate albino phenotype, along with corresponding intermediate *PDS* transcript and chlorophyll levels, reflecting the intentionally attenuated silencing strength rather than a progressive loss of activity (Figure 3). By contrast, conventional inverted-repeat RNAi transgenic constructs frequently undergo progressive loss of silencing due to transcriptional inactivation driven by DNA methylation of repetitive hairpin structures and associated promoters (Dong et al., 2011; Lu et al., 2015; Zhang et al., 2022). In comparison, the mirtron architecture lacks long inverted repeats and instead relies on a single short native intron, thereby reducing structural features known to trigger transgene methylation (Ruby et al., 2007; Westholm and Lai, 2011). Furthermore, mirtron sequences can be introduced into endogenous introns via homology-directed recombination, allowing precise integration and reducing transgene-associated silencing.

The durability of RNAi-based technologies in the field is challenged by pathogens that encode potent suppressors of host RNA silencing. Viral suppressors of RNA silencing, such as the *Tomato bushy stunt virus* P19 protein and the 2b suppressor from *Cucumber mosaic virus*, bind and sequester 21-nucleotide siRNA duplexes, thereby preventing their loading into the RISC (Voinnet, 2003; Lakatos et al., 2006). In addition to suppressing antiviral RNAi, these viral proteins can interfere with both native and artificial small RNA pathways. Transgenic expression of P19 in tomato, in the absence of viral infection, caused developmental abnormalities resembling viral disease symptoms, a phenotype directly linked to sequestration of endogenous miRNA/miRNA* duplexes, reduced microRNA activity, and accumulation of miRNA target transcripts. Thus, P19 suppresses not only antiviral siRNAs but also endogenous microRNA-mediated gene regulation. This counter-defense mechanism enables viruses to evade host RNAi and has been shown to compromise the efficacy of HIGS. In laboratory assays, co-expression of P19 or 2b abolished transgene-induced RNAi, preventing hairpin-mediated silencing of reporter genes in *Nicotiana benthamiana* (Voinnet, 2003; Lakatos et al., 2006). In crops that are frequently affected by viral infections, such as tomato, potato, cassava, and cucurbits, RNAi-based resistance traits have likewise been reported to be weakened in the presence of viral silencing suppressors, highlighting the vulnerability of RNAi and HIGS strategies under pathogen pressure (Dunoyer et al., 2010; Pumplin and Voinnet, 2013). The results of this study showed that mirtron-mediated *PDS* silencing was maintained in the presence of P19, whereas amiRNA-based silencing was frequently suppressed (Figure 5). Several unrelated viral silencing suppressors act through similar small RNA sequestration mechanisms (Voinnet, 2005; Ding and Voinnet, 2007). These findings suggest that mirtron-based silencing may provide enhanced robustness for HIGS applications, particularly under pathogen-rich field conditions.

Beyond stability and resistance to silencing suppressors, mirtron-based constructs can be used to target gene families. In the present study, a single *miR160*-based mirtron was shown to coordinately silence *StARF10* and *StARF17* in potato, resulting in characteristic developmental phenotypes such as dwarfism and pronounced leaf curvature, thereby demonstrating that mirtrons can efficiently target gene families and genetically redundant pathways (Figure 4;Kondhare et al., 2020). Consistent with these findings, studies in *Arabidopsis* have shown that *miR160* plays a key regulatory role during regeneration and is associated with impaired callus formation and reduced shoot regeneration capacity (Qiao et al., 2012; Qiao and Xiang, 2013; López-Ruiz et al., 2019). In line with these observations, only moderate *StARF10* and *StARF17* silencing was detected in 35S::GFP-MIRT_160_ transgenic potato lines, in contrast to the more pronounced silencing achieved for *PDS* in *Arabidopsis*. This difference likely reflects a regeneration bias, as transgenic lines exhibiting strong *miR160*-based mirtron silencing failed to regenerate. Together, these results highlight the modular nature of mirtron constructs, which enables flexible design strategies that permit either simultaneous targeting of multiple members of a gene family or selective silencing of individual genes.

From an applied perspective, the mirtron platform integrates the advantages of existing PTGS technologies, including tunability, reversibility, tissue specificity, and applicability to HIGS, while avoiding several limitations of conventional RNAi. This is particularly relevant for crop traits that benefit from tissue-specific modulation, such as fruit-restricted ripening (Gupta et al., 2013), vascular-specific HIGS (Alakonya et al., 2012), or localized modification of primary and secondary metabolism (Davuluri et al., 2005, (Sunilkumar et al., 2006; Wood et al., 2018). From a regulatory standpoint, mirtron produce only non-coding RNAs derived from intronic sequences. Following targeted integration of a mirtron via HDR and removal of T-DNA sequences, mirtron-containing plants resemble gene-edited products that lack any foreign sequence, including new promoters, selectable markers, and viral elements, as exemplified by a tomato product harboring an artificial mirtron that was recently exempted from USDA regulation (https://www.aphis.usda.gov/sites/default/files/25-191-01air-response.pdf).

In conclusion, the establishment of a functional mirtron pathway in plants represents both a conceptual advance in RNA silencing biology and a practical innovation for crop biotechnology. By coupling splicing to small RNA biogenesis, mirtron-mediated silencing provides a stable, flexible, and versatile platform that expands the toolkit for developing resilient and adaptable crops.

## Materials and Methods

### Plant material

Experiments were performed using *Arabidopsis thaliana* (ecotype Columbia-0, Col-0) and potato (*Solanum tuberosum* L. cv. Désirée). *Arabidopsis* plants were germinated on agar plates containing half-strength Murashige and Skoog medium (MS; Duchefa Biochemie, The Netherlands) under controlled growth conditions at 22 °C with a 16-h light/8-h dark photoperiod. For soil growth, both *Arabidopsis* and potato plants were transferred to a soil mixture consisting of 30% vermiculite, 30% peat, 20% tuff, and 20% perlite (Even Ari, Israel). Plants were grown in a temperature-controlled greenhouse under natural photoperiod conditions.

### Plasmid construction and mirtron design

All DNA constructs were synthesized by GenScript (Piscataway, NJ, USA) and cloned into the binary vector pPZP (Hajdukiewicz et al., 1994). Synthetic mirtrons were designed according to established principles of artificial mirtron architecture, in which splicing is defined by the 5′ splice donor site, branch point, and 3′ splice acceptor site (Salim et al., 2022; Kock et al., 2015) Mirtrons were engineered to target *Arabidopsis PHYTOENE DESATURASE 3* (*PDS3*, AT4G14210) or potato *StARF10* (PGSC0003DMT400020874) and *StARF17* (PGSC0003DMT400071449), using the seed sequence of *miR160* for the latter. All mirtron- or intron-containing sequences were embedded within the *GFP* coding sequence, inserted between nucleotides 13 and 14 of the *GFP* CDS. The primary construct, 35S::GFP-MIRT_PDS_ (construct 94), contains a synthetic mirtron harboring a seed sequence targeting *PDS3* (nucleotides 21–41 of the CDS) and is expressed under the cauliflower mosaic virus *35S* promoter and terminator. As a splicing control, 35S::GFP-INT was generated by inserting the 15th intron of *Arabidopsis* AT3G60950 into the same position within the *GFP* coding sequence. Constructs 9483, and 9482 were derived from construct 94. Construct 9483 contains an A-to-G substitution at position 23 downstream of the 5′ splice site. Construct 9482 (35S::GFP-MIRTPDS-NSC) carries an A-to-G substitution at position 13 downstream of the 5′ splice site. To modulate silencing strength, 35S::GFP-MIRT_PDS(A18T)_ was generated by introducing a single nucleotide substitution (A to T) at position 18 within the mirtron seed region. For multiplex silencing experiments in potato, 35S::GFP-MIRT_160_ was constructed by replacing the *PDS* seed sequence with the potato *miR160* seed sequence while maintaining the same mirtron architecture. To assess sensitivity to viral silencing suppression, mirtron and amiRNA constructs were expressed under the moderate constitutive *pCH1* promoter (1001 bp upstream of AT1G23490; (Zhou et al., 2023). pCH1::GFP-MIRT_PDS_ expresses the *PDS*-targeting mirtron under the *pCH1* promoter, while pCH1::amiRNA_PDS_ contains a previously described amiRNA targeting *PDS* (Tretter et al., 2008). For co-expression assays, an additional cassette encoding the *Tomato bushy stunt virus* silencing suppressor *P19*, driven by the 35S promoter and *NOS* terminator, was introduced into the corresponding pCH1::GFP-MIRT_PDS_ and pCH1::amiRNA_PDS_ constructs to generate pCH1::GFP-MIRT_PDS_+P19 and pCH1::amiRNA_PDS_+P19, respectively.

### Generation of transgenic plants

Electrocompetent *Agrobacterium tumefaciens* strain GV3101 was transformed by electroporation with 100 ng of plasmid DNA and used for plant transformation. Stable transformation of *Arabidopsis* (Col-0) was performed using the floral dip method (Clough and Bent, 1998). Transgenic *Arabidopsis* plants were selected on half-strength MS medium containing 1% sucrose and 50 mg L^-1^ kanamycin. Potato transformation was carried out using an *Agrobacterium*-mediated protocol adapted from Ginzberg et al. (2012). Fully expanded sterile leaves were excised into discs and immersed for 5–10 min in an *Agrobacterium* suspension prepared in inoculation buffer (MS liquid medium) at OD_600_ = 0.5. Explants were co-cultivated for 48 h in the dark on MS medium supplemented with 3% sucrose (Duchefa, Haarlem, The Netherlands) and 200 µM acetosyringone (Sigma-Aldrich, St. Louis, MO, USA). Following co-cultivation, explants were transferred to callus induction medium consisting of MS salts, 3% sucrose, 0.1 mg L⁻¹ 6-benzylaminopurine (BAP), and 5 mg L^-1^ naphthalene acetic acid (NAA), supplemented with 500 mg L^-1^ cefotaxime and 50 mg L^-1^ kanamycin (Duchefa unless otherwise stated), and maintained at 24 °C under a 16 h light/8 h dark photoperiod for 10 days. For shoot development and selection, explants were transferred to MS-based medium containing 3% sucrose, 2 mg L^-1^ zeatin riboside, 0.02 mg L^-1^ gibberellic acid (GA_3_), 0.02 mg L^-1^ NAA, 500 mg L^-1^ cefotaxime, and 50 mg L^-1^kanamycin. Emerging shoots were excised after approximately six weeks and transferred to a rooting medium consisting of MS salts, 3% sucrose, 500 mg L^-1^ cefotaxime, and 50 mg L^-1^ kanamycin. Rooted plantlets were transplanted to soil, acclimated under controlled conditions for 10 days, and transferred to the greenhouse. Transgenic plants were confirmed by PCR analysis of genomic DNA (Supplementary Figure 2), using primers listed in Table S1.

### Chlorophyll Content Measurement

Approximately 80 mg fresh weight of leaf tissue was harvested 19 days after germination, flash-frozen in liquid nitrogen, and extracted in 1 mL of 90% (v/v) aqueous acetone. Samples were centrifuged at 8,000 × g for 10 min at 4 °C, and chlorophyll content was quantified as described by (Jeffrey and Humphrey, 1975). Absorbance was measured in 96-well plates using a spectrophotometer (INFINITE 200 PRO Reader; Tecan, Männedorf, Switzerland) at 630, 647, 664, and 750 nm, with 750 nm used for background correction. Chlorophyll concentrations (µg mL^-1^) were calculated as: chlorophyll *a* = 11.85A_664_ − 1.54A_647_ + 0.08A_630_; chlorophyll *b* = 5.43A_664_ + 21.03A_647_ − 2.66A_630_, and normalized to fresh weight. Four biological replicates, each measured in three technical replicates, were analyzed per treatment.

### Fluorescence microscopy

All fluorescence imaging was performed using a Magnus FL inverted microscope (Magnus, Uttar Pradesh, India). GFP fluorescence was visualized using an excitation wavelength of 470 nm, and emission was detected using a 505 nm cutoff filter.

### RNA extraction and quantitative real-time PCR

Total RNA was isolated from harvested plant tissues using the LogSpin method (Yaffe et al., 2012). Samples were homogenized in 8M guanidine hydrochloride (Duchefa Biochemie, The Netherlands) using a Bullet Blender Storm Pro (Next Advance, Inc., Troy, NY) and mixed with 96% ethanol prior to loading onto plasmid DNA purification columns (RBC Bioscience, New Taipei City, Taiwan). Columns were washed twice with 3 M sodium acetate BDH Chemicals (VWR International, Radnor, PA, USA) and twice with 75% ethanol, and RNA was eluted with DEPC-treated water (Biological Industries, Beit Haemek, Israel) preheated to 65 °C. Residual genomic DNA was removed by treatment with RQ1 DNase (Promega, Madison, WI). First-strand cDNA was synthesized from 1 μg total RNA using Maxima First Strand cDNA Synthesis Kit for RT-qPCR (ThermoFisher Scientific, Waltham, MA, USA) and diluted 1:5 prior to analysis. Quantitative real-time PCR was performed using SYBR Green master mix (Thermo Fisher Scientific) on a CFX96 Real-Time PCR System (Bio-Rad Laboratories, Inc., Hercules, CA, USA). Cycling conditions consisted of an initial denaturation at 95 °C for 3 min followed by 40 cycles of 95 °C for 10 s, 60 °C for 20 s, and 72 °C for 15 s. Data were analyzed using CFX Maestro software v2.3 (Bio-Rad Laboratories, Inc., Hercules, CA, USA). Transcript levels were normalized to *the Arabidopsis TUBULIN BETA CHAIN 2 gene (AT5G62690; TUB2) or the* potato *EUKARYOTIC TRANSLATION INITIATION FACTOR 3 SUBUNIT E* (*ELF3e*). Primer sequences are provided in Table S1. Relative expression levels were calculated using ΔΔCt.

## Data Availability Statement

All data supporting the findings of this study are available within the article and its Supporting Information.

## Author Contributions

N.L. and K.S. conceived the study; N.L. performed the experiments with assistance from S.N.; B.J.I. generated the transgenic potato lines and conducted and analyzed expression studies; B.S-R. contributed to data interpretation and critical conceptual input; N.L. and B.S-R. wrote the manuscript with input from B.J.I. and K.S.

## Acknowledgements

We thank Dr. Gusui Wu for his critical review of the manuscript and for his valuable, constructive suggestions. Generative AI (ChatGPT, GPT-4 family model, February 2026 version; OpenAI) was used solely for language editing and clarity improvement. All scientific content and interpretations were generated and verified by the authors.

## References

Alakonya, A., Kumar, R., Koenig, D., Kimura, S., Townsley, B., Runo, S., Garces, H. M., Kang, J., Yanez, A., & David-Schwartz, R. (2012). Interspecific RNA interference of SHOOT MERISTEMLESS-like disrupts Cuscuta pentagona plant parasitism. The Plant Cell, 24(7), 3153–3166.

Bramlett, M., Plaetinck, G., & Maienfisch, P. (2020). RNA-based biocontrols—a new paradigm in crop protection. Engineering, 6(5), 522–527.

Brown, J. W. S., Marshall, D. F., & Echeverria, M. (2008). Intronic noncoding RNAs and splicing. Trends in Plant Science, 13(7), 335–342.

Cardi, T., Murovec, J., Bakhsh, A., Boniecka, J., Bruegmann, T., Bull, S. E., Eeckhaut, T., Fladung, M., Galovic, V., & Linkiewicz, A. (2023). CRISPR/Cas-mediated plant genome editing: outstanding challenges a decade after implementation. Trends in Plant Science, 28(10), 1144–1165.

Charles, H., Godfray, J., Beddington, J. R., Crute, I. R., Haddad, L., Lawrence, D., Muir, J. F., Pretty, J., Robinson, S., Thomas, S. M., & Toulmin, C. (n.d.). Food Security: The Challenge of Feeding 9 Billion People. Retrieved www.sciencemag.org

Clough, S. J., & Bent, A. F. (1998). Floral dip: a simplified method for Agrobacterium-mediated transformation of Arabidopsis thaliana. The Plant Journal, 16(6), 735–743.

Davuluri, G. R., Van Tuinen, A., Fraser, P. D., Manfredonia, A., Newman, R., Burgess, D., Brummell, D. A., King, S. R., Palys, J., & Uhlig, J. (2005). Fruit-specific RNAi-mediated suppression of DET1 enhances carotenoid and flavonoid content in tomatoes. Nature Biotechnology, 23(7), 890–895.

Ding, S.-W., & Voinnet, O. (2007). Antiviral immunity directed by small RNAs. Cell, 130(3), 413–426.

Dunoyer, P., Brosnan, C. A., Schott, G., Wang, Y., Jay, F., Alioua, A., Himber, C., & Voinnet, O. (2010). Retracted: An endogenous, systemic RNAi pathway in plants. The EMBO Journal, 29(10), 1699–1712.

F de Felippes, F., McHale, M., Doran, R. L., Roden, S., Eamens, A. L., Finnegan, E. J., & Waterhouse, P. M. (2020). The key role of terminators on the expression and post-transcriptional gene silencing of transgenes. The Plant Journal, 104(1), 96–112.

Feng, Z., Li, X., Fan, B., Zhu, C., & Chen, Z. (2022). Maximizing the production of recombinant proteins in plants: from transcription to protein stability. International Journal of Molecular Sciences, 23(21), 13516.

Formey, D., Sallet, E., Lelandais-Brière, C., Ben, C., Bustos-Sanmamed, P., Niebel, A., Frugier, F., Combier, J. P., Debellé, F., Hartmann, C., Poulain, J., Gavory, F., Wincker, P., Roux, C., Gentzbittel, L., Gouzy, J., & Crespi, M. (2014). The small RNA diversity from Medicago truncatula roots under biotic interactions evidences the environmental plasticity of the miRNAome. Genome Biology, 15(9). 10.1186/s13059-014-0457-4

Ginzberg, I., Thippeswamy, M., Fogelman, E., Demirel, U., Mweetwa, A. M., Tokuhisa, J., & Veilleux, R. E. (2012). Induction of potato steroidal glycoalkaloid biosynthetic pathway by overexpression of cDNA encoding primary metabolism HMG-CoA reductase and squalene synthase. Planta, 235(6), 1341–1353.

Guo, Q., Liu, Q., A. Smith, N., Liang, G., & Wang, M.-B. (2016). RNA silencing in plants: mechanisms, technologies and applications in horticultural crops. Current Genomics, 17(6), 476–489.

Gupta, A., Pal, R. K., & Rajam, M. V. (2013). Delayed ripening and improved fruit processing quality in tomato by RNAi-mediated silencing of three homologs of 1-aminopropane-1-carboxylate synthase gene. Journal of Plant Physiology, 170(11), 987–995.

Hajdukiewicz, P., Svab, Z., & Maliga, P. (1994). The small, versatile pPZP family of Agrobacterium binary vectors for plant transformation. Plant Molecular Biology, 25(6), 989–994.

He, S., & Krainer, K. M. C. (2021). The inequity of biotechnological impact. Molecular Plant, 14(1), 1–2.

Hickey, L. T., N. Hafeez, A., Robinson, H., Jackson, S. A., Leal-Bertioli, S. C. M., Tester, M., Gao, C., Godwin, I. D., Hayes, B. J., & Wulff, B. B. H. (2019). Breeding crops to feed 10 billion. In Nature Biotechnology (Vol. 37, Number 7, pp. 744–754). Nature Research. 10.1038/s41587-019-0152-9

Jeffrey, S. W. t, & Humphrey, G. F. (1975). New spectrophotometric equations for determining chlorophylls a, b, c1 and c2 in higher plants, algae and natural phytoplankton. Biochemie Und Physiologie Der Pflanzen, 167(2), 191–194.

Joshi, P. K., Gupta, D., Nandal, U. K., Khan, Y., Mukherjee, S. K., & Sanan-Mishra, N. (2012). Identification of mirtrons in rice using MirtronPred: A tool for predicting plant mirtrons. Genomics, 99(6), 370–375. 10.1016/j.ygeno.2012.04.002

Kamthan, A., Chaudhuri, A., Kamthan, M., & Datta, A. (2015). Small RNAs in plants: recent development and application for crop improvement. Frontiers in Plant Science, 6, 208.

Kock, K. H., Kong, K. W., Hoon, S., & Seow, Y. (2015). Functional VEGFA knockdown with artificial 3′-tailed mirtrons defined by 5′ splice site and branch point. Nucleic Acids Research, 43(13), 6568–6578.

Kondhare, K. R., Natarajan, B., & Banerjee, A. K. (2020). Molecular signals that govern tuber development in potato. International Journal of Developmental Biology, 64(1-2–3), 133–140.

Koo, H., Jung, M., Lee, S., Go, S., & Kim, Y.-M. (2025). Identification and application of promoters and terminators for plant synthetic biology. Molecules and Cells, 100273.

Kumar, M., Chaudhary, V., Yadav, M. K., Chauhan, C., Kumar, R., Singh, D., & Teotia, S. (2025). RNAi: a potent biotechnological tool for improvement of ornamental crops. Plant Molecular Biology Reporter, 43(1), 23–40.

Lakatos, L., Csorba, T., Pantaleo, V., Chapman, E. J., Carrington, J. C., Liu, Y.-P., Dolja, V. V, Calvino, L. F., López-Moya, J. J., & Burgyán, J. (2006). Small RNA binding is a common strategy to suppress RNA silencing by several viral suppressors. The EMBO Journal, 25(12), 2768.

Lopez-Gomollon, S., & Baulcombe, D. C. (2022). Roles of RNA silencing in viral and non-viral plant immunity and in the crosstalk between disease resistance systems. Nature Reviews Molecular Cell Biology, 23(10), 645–662.

López-Ruiz, B. A., Juárez-González, V. T., Sandoval-Zapotitla, E., & Dinkova, T. D. (2019). Development-related miRNA expression and target regulation during staggered in vitro plant regeneration of tuxpeño VS-535 maize cultivar. International Journal of Molecular Sciences, 20(9), 2079.

Lu, S., Yin, X., Spollen, W., Zhang, N., Xu, D., Schoelz, J., Bilyeu, K., & Zhang, Z. J. (2015). Analysis of the siRNA-mediated gene silencing process targeting three homologous genes controlling soybean seed oil quality. PLoS One, 10(6), e0129010.

Matzke, A. J. M., & Matzke, M. A. (1998). Position effects and epigenetic silencing of plant transgenes. Current Opinion in Plant Biology, 1(2), 142–148.

Pixley, K. V, Falck-Zepeda, J. B., Paarlberg, R. L., Phillips, P. W. B., Slamet-Loedin, I. H., Dhugga, K. S., Campos, H., & Gutterson, N. (2022). Genome-edited crops for improved food security of smallholder farmers. Nature Genetics, 54(4), 364–367.

Pumplin, N., & Voinnet, O. (2013). RNA silencing suppression by plant pathogens: defence, counter-defence and counter-counter-defence. Nature Reviews Microbiology, 11(11), 745–760.

Qiao, M., & Xiang, F. (2013). A set of Arabidopsis thaliana miRNAs involve shoot regeneration in vitro. Plant Signaling & Behavior, 8(3), e23479.

Qiao, M., Zhao, Z., Song, Y., Liu, Z., Cao, L., Yu, Y., Li, S., & Xiang, F. (2012). Proper regeneration from in vitro cultured Arabidopsis thaliana requires the microRNA-directed action of an auxin response factor. The Plant Journal, 71(1), 14–22.

Roca Paixao, J. F., & Déléris, A. (2025). Epigenetic control of T-DNA during transgenesis and pathogenesis. Plant Physiology, 197(1), kiae583.

Ruby, J. G., Jan, C. H., & Bartel, D. P. (2007). Intronic microRNA precursors that bypass Drosha processing. Nature, 448(7149), 83–86.

Salim, U., Kumar, A., Kulshreshtha, R., & Vivekanandan, P. (2022). Biogenesis, characterization, and functions of mirtrons. Wiley Interdisciplinary Reviews: RNA, 13(1), e1680.

Saurabh, S., Vidyarthi, A. S., & Prasad, D. (2014). RNA interference: concept to reality in crop improvement. Planta, 239(3), 543–564.

Shapulatov, U., van Hoogdalem, M., Schreuder, M., Bouwmeester, H., Abdurakhmonov, I. Y., & van der Krol, A. R. (2018). Functional intron-derived miRNAs and host-gene expression in plants. Plant Methods, 14(1), 83.

Shomron, N., & Levy, C. (2009). MicroRNA-biogenesis and pre-mRNA splicing crosstalk. BioMed Research International, 2009(1), 594678.

Sunilkumar, G., Campbell, L. M., Puckhaber, L., Stipanovic, R. D., & Rathore, K. S. (2006). Engineering cottonseed for use in human nutrition by tissue-specific reduction of toxic gossypol. Proceedings of the National Academy of Sciences, 103(48), 18054–18059.

Tretter, E. M., Alvarez, J. P., Eshed, Y., & Bowman, J. L. (2008). Activity range of Arabidopsis small RNAs derived from different biogenesis pathways. Plant Physiology, 147(1), 58–62.

Turnbull, C., Lillemo, M., & Hvoslef-Eide, T. A. K. (2021). Global regulation of genetically modified crops amid the gene edited crop boom–a review. Frontiers in Plant Science, 12, 630396.

Ullah, I., Kamel, E. A. R., Shah, S. T., Basit, A., Mohamed, H. I., & Sajid, M. (2022). Application of RNAi technology: a novel approach to navigate abiotic stresses. Molecular Biology Reports, 49(11), 10975–10993.

Vaucheret, H., & Voinnet, O. (2024). The plant siRNA landscape. The Plant Cell, 36(2), 246–275.

Voinnet, O. (2003). RNA silencing bridging the gaps in wheat extracts. Trends in Plant Science, 8(7), 307–309.

Voinnet, O. (2005). Induction and suppression of RNA silencing: insights from viral infections. Nature Reviews Genetics, 6(3), 206–220.

Voinnet, O. (2009). Origin, biogenesis, and activity of plant microRNAs. Cell, 136(4), 669–687.

Westholm, J. O., & Lai, E. C. (2011). Mirtrons: microRNA biogenesis via splicing. Biochimie, 93(11), 1897–1904.

Wood, C. C., Okada, S., Taylor, M. C., Menon, A., Mathew, A., Cullerne, D., Stephen, S. J., Allen, R. S., Zhou, X., & Liu, Q. (2018). Seed-specific RNAi in safflower generates a superhigh oleic oil with extended oxidative stability. Plant Biotechnology Journal, 16(10), 1788–1796.

Yaffe, H., Buxdorf, K., Shapira, I., Ein-Gedi, S., Moyal-Ben Zvi, M., Fridman, E., Moshelion, M., & Levy, M. (2012). LogSpin: a simple, economical and fast method for RNA isolation from infected or healthy plants and other eukaryotic tissues. BMC Research Notes, 5(1), 45.

Zhang, D., Zhong, C., Smith, N. A., de Feyter, R., Greaves, I. K., Swain, S. M., Zhang, R., & Wang, M.-B. (2022). Nucleotide mismatches prevent intrinsic self-silencing of hpRNA transgenes to enhance RNAi stability in plants. Nature Communications, 13(1), 3926.

Zhou, A., Kirkpatrick, L. D., Ornelas, I. J., Washington, L. J., Hummel, N. F. C., Gee, C. W., Tang, S. N., Barnum, C. R., Scheller, H. V, & Shih, P. M. (2023). A suite of constitutive promoters for tuning gene expression in plants. ACS Synthetic Biology, 12(5), 1533–1545.

