## Supplementary Figures and Table for "A Synthetic Mirtron Platform Enables Stable and Robust Splicing-Dependent Gene Silencing in Plants"

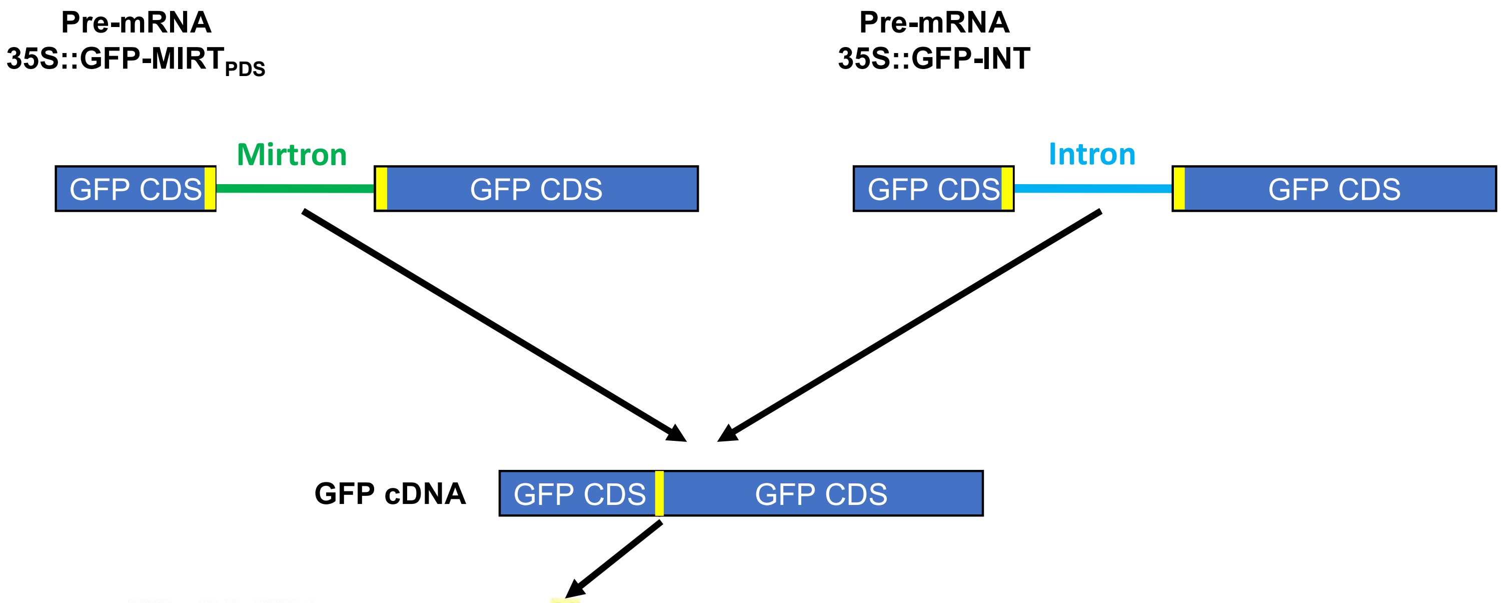


```
35S::GFP-MIRTPDS    ATGGTGAGCAAAGGTGAGGAGCTGTTCACCGGGGTGGTGCCCATCCTGGTCGAGCTGGAC    60
35S::GFP-INT        ATGGTGAGCAAAGGTGAGGAGCTGTTCACCGGGGTGGTGCCCATCCTGGTCGAGCTGGAC    60
                    ************************************************************

35S::GFP-MIRTPDS    GGCGACGTAAACGGCCACAAGTTCAGCGTGTCCGGCGAGGGCGAGGGCGATGCCACCTAC    120
35S::GFP-INT        GGCGACGTAAACGGCCACAAGTTCAGCGTGTCCGGCGAGGGCGAGGGCGATGCCACCTAC    120
                    ************************************************************

35S::GFP-MIRTPDS    GGCAAGCTGACCCTGAAGTTCATCTGCACCACCGGCAAGCTGCCCGTGCCCTGGCCCACC    180
35S::GFP-INT        GGCAAGCTGACCCTGAAGTTCATCTGCACCACCGGCAAGCTGCCCGTGCCCTGGCCCACC    180
                    ************************************************************

35S::GFP-MIRTPDS    CTCGTGACCACCTTCACCTACGGCGTGCAGTGCTTCAGCCGCTACCCCGACCACATGAAG    240
35S::GFP-INT        CTCGTGACCACCTTCACCTACGGCGTGCAGTGCTTCAGCCGCTACCCCGACCACATGAAG    240
                    ************************************************************

35S::GFP-MIRTPDS    CAGCACGACTTCTTCAAGTCCGCC    264
35S::GFP-INT        CAGCACGACTTCTTCAAGTCCGCC    264
                    ************************
```

**Supplementary Figure 1.** Sequence verification of GFP-based mirtron and intron constructs. Schematic representation of the GFP reporter constructs used in this study. The synthetic mirtron (left, in green) or control intron (right, in blue) was inserted within the *GFP* coding sequence. Canonical splice donor (GT) and acceptor (AG) sites are highlighted in yellow.
Bottom panel: Alignment of cDNA sequencing results of 35S::GFP-MIRT$_{PDS}$ and 35S::GFP-INT constructs across the insertion region. The guanine nucleotide remaining at the exon–exon junction following splicing is highlighted in yellow. Asterisks denote identical nucleotides between constructs.

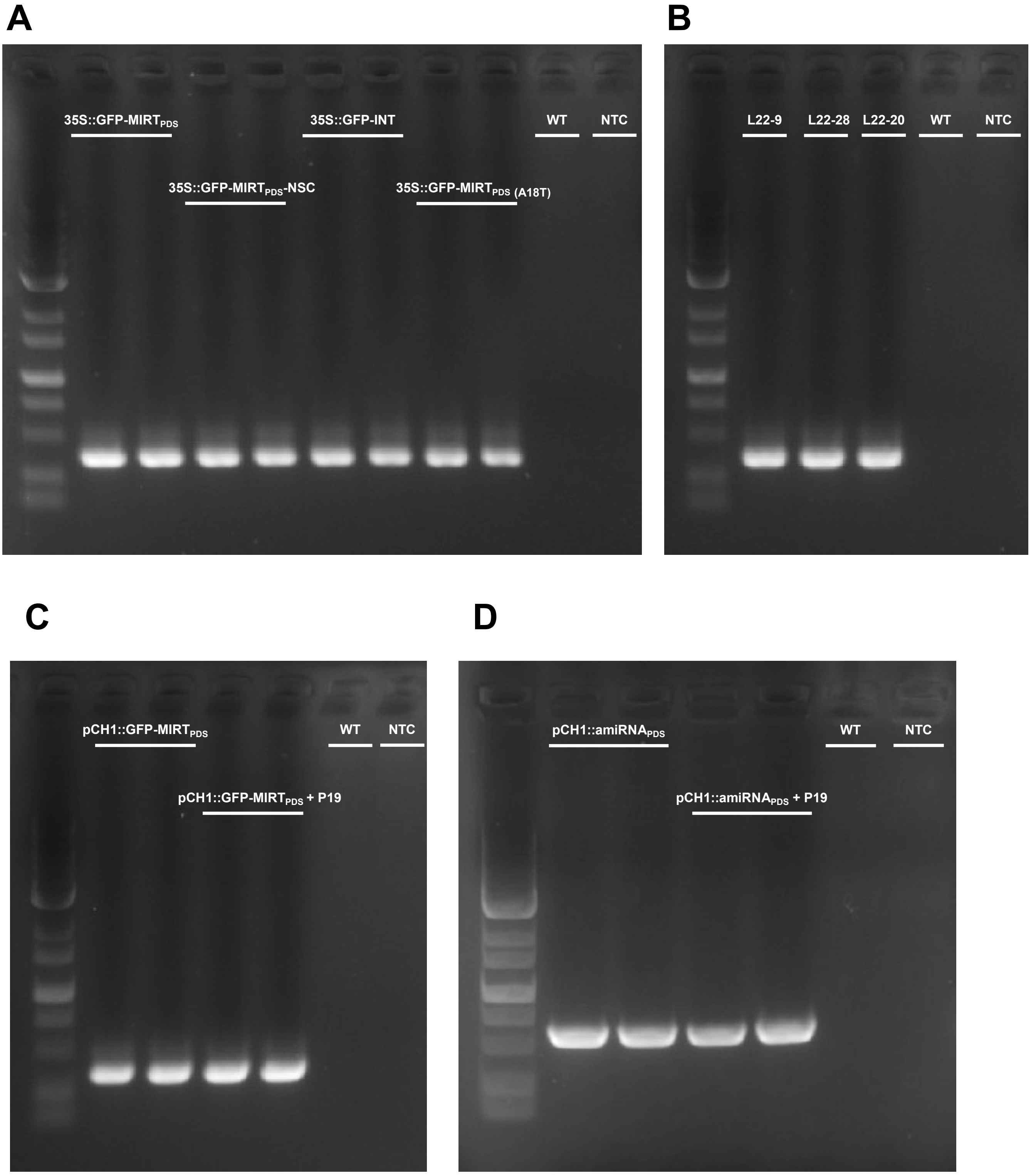


**Supplemental Figure 2.** Molecular validation of transgenic lines. (A) PCR confirmation of transgenic lines carrying the constructs 35S::GFP-MIRT$_{PDS}$, 35S::GFP-MIRT$_{PDS}$-NSC, 35S::GFP-INT, and 35S::GFP-MIRT$_{PDS\ (A75T).}$ Amplification was performed using primers flanking the mirtron/intron region (T-DNA1; see Table S1). (B) PCR confirmation of lines harboring the 35S::GFP-MIRT$_{160}$ construct. Primers flanking the mirtron region were used for amplification (T-DNA1; Table S1). (C) PCR confirmation of lines transformed with pCH1::GFP-MIRT$_{PDS}$ and pCH1::GFP-MIRT$_{PDS}$+P19. Amplification was performed using primers flanking the mirtron sequence (T-DNA1; Table S1). (D) PCR confirmation of lines carrying pCH1::amiRNA$_{PDS}$ and pCH1::amiRNA$_{PDS}$+P19 constructs. Amplification was performed using primers flanking the amiRNA sequence (T-DNA2; Table S1). All PCR reactions were carried out using genomic DNA as a template.

**Table S1**
**Primers list**

| Gene | Forward primer (5'-3') | Reverse primer (5'-3') | Experiment |
|---|---|---|---|
| **PDS** | GGTATTTGGGCTATTTTGCG | CTCCCTGCTTTTCCATCCA | Q-PCR |
| **TUB2** | AAACTCACTACCCCCAGCTTTG | CACCAGACATAGTAGCAGAAATCAAGT | Q-PCR |
| **ARF10** | GTCCAGCAGTCCTTTCTGTTGTTT | GCTGCAACACGCTGGAAACTT | Q-PCR |
| **ARF17** | TGAAGGCACAAGCGGTAGAG | CTCCGTTTCCACAGCCATCT | Q-PCR |
| **EIF3e** | GGAGCACAGGAGAAGATGAAGGAG | CGTTGGTGAATGCGGCAGTAGG | Q-PCR |
| **T-DNA1** | GAAAATTTTCACCATTTACGAACGATAGTG | GGCGGACTTGAAGAAGTCGT | Transgene confirmation – Fig. S3 A-C |
| **T-DNA2** | GAAAATTTTCACCATTTACGAACGATAGTG | GAACCACCCATAATACCCATAATAGCT | Transgene confirmation – Fig. S3 D |
| **GFP** | ATGGTGAGCAAAGGTGAGGAGC | GGCGGACTTGAAGAAGTCGT | cDNA sequencing |
